# Regulatory Dynamics of Sch9 in Response to Cytosolic Acidification: From Spatial Reconfiguration to Cellular Adaptation to Stresses

**DOI:** 10.1101/2024.03.12.584575

**Authors:** Rui Fujii, Eigo Takeda, Eisuke Itakura, Akira Matsuura

**Affiliations:** Department of Biology, Graduate School of Science and Engineering, Chiba University, Chiba 263-8522, Japan; Research Center for Cell Biology, Institute of Innovative Research, Tokyo Institute of Technology, Yokohama 226-8503, Japan; Department of Biology, Graduate School of Science, Chiba University, Chiba 263-8522, Japan

## Abstract

The regulation of cellular metabolism in response to intracellular and extracellular conditions is critical for cell survival. In *Saccharomyces cerevisiae*, Sch9 is a well-established substrate of the target of rapamycin complex 1 (TORC1) and regulates metabolic pathways and stress responses. Sch9 is enriched on the vacuolar membrane through binding to PI(3,5)P_2_, and this localization is essential for TORC1-dependent phosphorylation. Previous studies have demonstrated that glucose starvation and oxidative stress cause the dissociation of Sch9 from the vacuolar membrane. However, the underlying mechanism and physiological significance of the change in Sch9 localization still require elucidation. In this study, we demonstrated that cytosolic pH is a regulator of Sch9 localization. We observed that multiple stress conditions that induce cytosolic acidification consistently led to the detachment of Sch9 from the vacuolar membrane. Furthermore, we confirmed that the affinity between Sch9 and PI(3,5)P_2_ is pH-dependent *in vitro*. This pH-dependent localization switch of Sch9 is linked to selective regulation of the TORC1–Sch9 pathway. Impairment of the dissociation of Sch9 from the vacuolar membrane in response to cytosolic acidification resulted in deficient induction of the expression of the stress response gene and delayed the adaptive response to acetic acid stress. These findings indicate that the appropriate control of Sch9 localization is essential for metabolic reprogramming.

## Introduction

Sch9 is an AGC family protein kinase that is among the most evolutionarily conserved kinases (Urban et al., 2007). It is involved in various cellular processes, including growth (Jorgensen et al., 2002; Toda et al., 1988), metabolism (Swinnen et al., 2014), stress response (Oh et al., 2018; Pascual-Ahuir and Proft, 2007; Pedruzzi et al., 2003; Yorimitsu et al., 2007), and lifespan (Fabrizio et al., 2001; Kaeberlein et al., 2005). Similar to typical AGC family kinases, its activation requires the phosphorylation of its regulatory motifs. The C-terminus of Sch9, which contains hydrophobic motifs, is phosphorylated by the target of rapamycin complex 1 (TORC1), a highly conserved protein kinase complex (Urban et al., 2007). In addition, Sch9 possesses an additional phosphorylation site in the activation loop that is phosphorylated by Pkh1 and Pkh2, which are orthologs of mammalian PDK1 (Liu et al., 2005; Roelants et al., 2004; Urban et al., 2007; Voordeckers et al., 2011).

One significant function of Sch9 is its promotion of ribosome biogenesis. Once activated, Sch9 directly phosphorylates and inhibits Dot6, Tod6, and Stb3, repressors of ribosome biogenesis (Ribi) and ribosomal protein (RP) genes (Huber et al., 2011). In addition, it phosphorylates and inhibits Maf1, a negative regulator of RNA polymerase III, thereby promoting the transcription of 5S rRNA and tRNAs (Huber et al., 2009). It also plays a significant role in stress responses. In nutrient-rich conditions, activated Sch9 phosphorylates and sequesters the Greatwall protein kinase Rim15 away from the nucleus (Pedruzzi et al., 2003; Wanke et al., 2005). By contrast, nutrient-poor conditions relieve the inhibitory phosphorylation of Sch9 on Rim15; then, Rim15 is translocated into the nucleus, where it induces the expression of stress-response element (STRE)-regulated and post-diauxic shift (PDS) element-regulated genes through Msn2/4 and Gis1, respectively (Pedruzzi et al., 2003; Roosen et al., 2005).

Previous studies have demonstrated that Sch9 is enriched on the vacuolar membrane (Jorgensen et al., 2004; Urban et al., 2007). The interaction between the N-terminal domain of Sch9 and the phospholipid phosphatidylinositol 3,5-bisphosphate (PI(3,5)P_2_) facilitates the localization of Sch9 to the vacuolar membrane (Jin et al., 2014; Novarina et al., 2021). The disruption of genes involved in PI(3,5)P_2_ synthesis, such as *VAC7* and *VAC14*, or truncation of the N-terminal domain of Sch9, results in impairment of the association between Sch9 and the vacuolar membrane. Loss of localization to the vacuolar membrane leads to the dephosphorylation of Sch9, emphasizing the essential role of vacuolar membrane localization in TORC1-dependent phosphorylation of Sch9 (Novarina et al., 2021). Under physiological conditions, glucose starvation and oxidative stress cause the dissociation of Sch9 from the vacuolar membrane (Jorgensen et al., 2004; Takeda et al., 2017; Wilms et al., 2017). Notably, this change contributes to the selective inhibition of the Sch9 branch within the TORC1 signaling pathway (Takeda et al., 2017). However, the mechanism by which the localization change in Sch9 is triggered and its physiological significance in stress responses remain unclear.

In this study, we reveal that Sch9 dissociates from the vacuolar membrane in response to a drop in cytosolic pH. This pH-dependent change in Sch9 localization contributes to the selective inactivation of the TORC1–Sch9 pathway and regulation of PDS gene expression. We further demonstrate that the dissociation of Sch9 from the vacuolar membrane promotes adaptation to acetic acid. Thus, we propose that cytosolic pH links appropriate metabolic regulation mediated by the control of Sch9 localization and activity.

## Results

### Sch9 dissociates from the vacuolar membrane in response to cytosolic acidification

As noted above, Sch9 is abundant on the vacuolar membrane but dissociates from it under glucose starvation and oxidative stress (Jorgensen et al., 2004; Takeda et al., 2017; Wilms et al., 2017). To explore whether other conditions affect its localization, we employed fluorescence microscopy to observe the subcellular localization of green fluorescent protein (GFP)–Sch9 under various conditions. We identified three distinct patterns of change in Sch9 dynamics. The initial pattern is the dissociation of Sch9 from the vacuolar membrane.

Consistent with previous findings, this pattern was observed under glucose deprivation and oxidative stress conditions (Figure 1A, B). Acetic acid treatment also caused it (Figure 1C). The second pattern involved an increase in the fraction of Sch9 on the vacuolar membrane, which was observed upon hyperosmotic stress, DNA damage, and α-factor treatment (Figure 1D–F). The third pattern revealed no significant change in localization compared to the untreated condition, as observed under rapamycin treatment (Figure 1G). Notably, almost all conditions tested in this study induced the dephosphorylation of Sch9, indicating that its phosphorylation state does not affect its localization. This hypothesis is supported by evidence demonstrating that phosphomimetic and non-phosphorylatable mutants of Sch9 exhibited localization patterns similar to that of wild-type (WT) Sch9 (Urban et al., 2007). Because the dissociation of Sch9 from the vacuolar membrane could be involved in its inactivation (Takeda et al., 2017; Urban et al., 2007), we focused on the first pattern hereafter.

**Figure 1.**
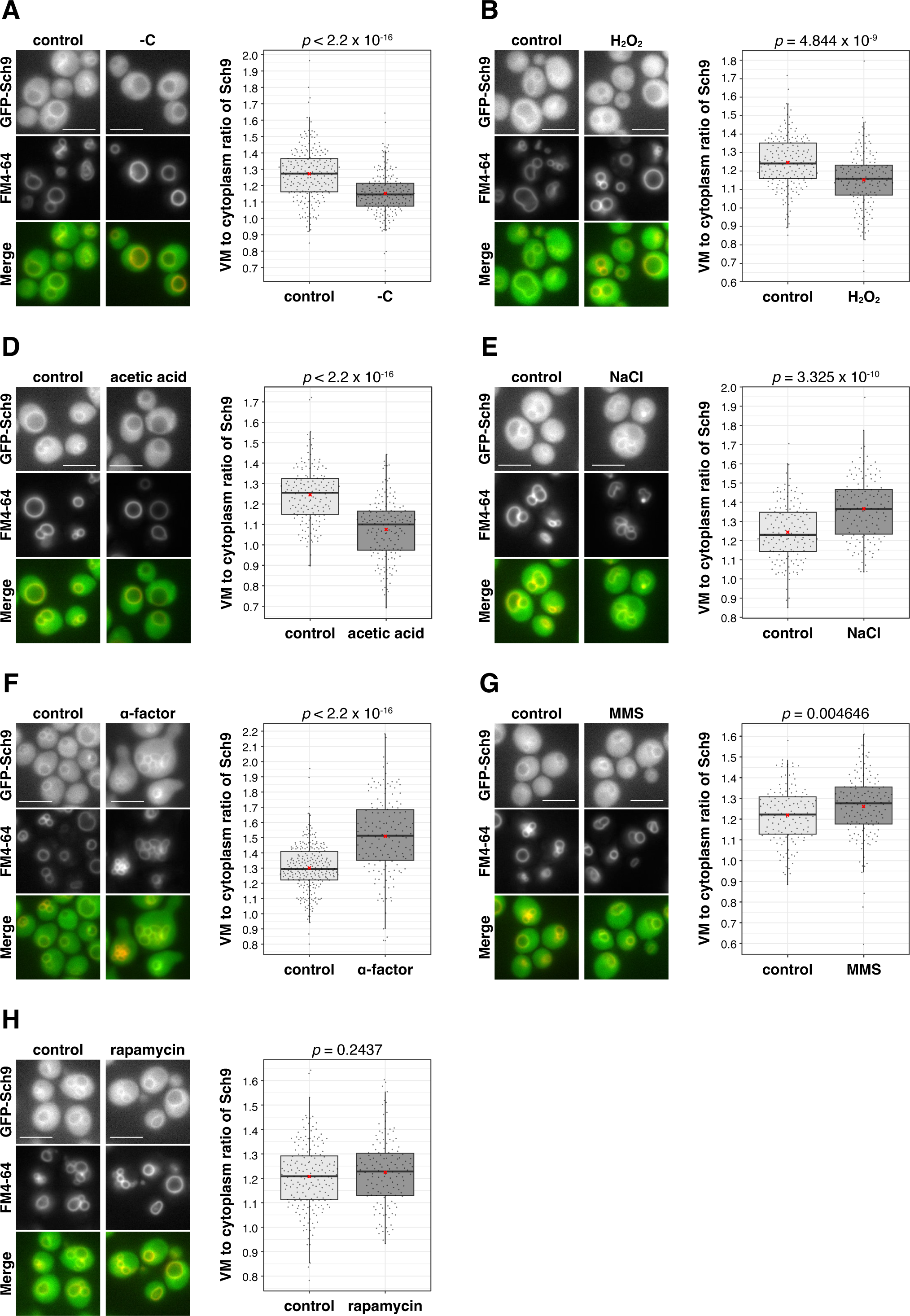
Sch9 localization under various conditions. **(A)**–**(G)** Left: Fluorescent images of GFP–Sch9 and a vacuole labeled with FM4–64 under control and stress treatment conditions. Right: Quantitative analysis of the ratio of Sch9 fluorescence intensity at vacuolar membranes to that in the cytoplasm. Exponentially growing cells were subjected to various stress conditions: glucose starvation for 30 min (A), oxidative stress (3 mM H_2_O_2_) for 30 min (B), acetic acid stress (35 mM acetic acid) for 30 min (C), hyperosmotic stress (0.8 M NaCl) for 30 min (D), treatment with 20 μg/mL α-factor for 2 h (E), DNA damage (0.03% MMS) for 30 min (F), or treatment with 100 ng/mL rapamycin for 30 min (G). Control cells were incubated in SDCU medium at 30°C for a duration corresponding to that of the stress treatment. Scale bars = 5 μm. In the boxplot, the bottom and top of the box represent the first (Q1) and third (Q3) quartiles, indicating the 25th and 75th percentiles, respectively. The band inside the box indicates the median (50th percentile). The whiskers extend from the box to the lowest and highest values within 1.5 × the interquartile range (IQR) from the first and third quartiles, respectively. Red crosses indicate mean values. Significance was evaluated using the Brunner–Munzel test.

The level of PI(3,5)P_2_, a scaffold of Sch9 on the vacuolar membrane, affects the localization of Sch9 (Jin et al., 2014). However, a previous study showed that Sch9 dissociates from the vacuolar membrane although the PI(3,5)P_2_ level does not decrease under oxidative stress (Takeda et al., 2017). Therefore, we assumed that factors other than the PI(3,5)P_2_ level affect Sch9 localization. The conditions leading to Sch9 dissociation from the vacuolar membrane induce a decrease in cytosolic pH (Dechant et al., 2010; Dodd and Kralj, 2017; Pampulha and Loureiro-Dias, 1989; Triandafillou et al., 2020). Indeed, glucose deprivation, oxidative stress, and acetic acid treatment reduced cytosolic pH, from ∼7.5 to ∼6.3, whereas other stresses did not, as monitored by pHluroin2, a GFP-based pH sensor (Figure S1A). Therefore, we hypothesized that a drop in cytosolic pH could mediate the dissociation of Sch9 from the vacuolar membrane. To test this hypothesis, we examined its subcellular localization after modulating cytosolic pH by several different methods.

First, we treated cells with weak acids, including sorbic and benzoic acids, which lowered the cytosolic pH from ∼7.5 to ∼6.5, resulting in less localization (Figures 2A, B, and S1B). Next, ebselen, a drug that inhibits the P-type ATPase Pma1 and rapidly lowers cytosolic pH (Chan et al., 2007), was used, which also reduced localization (Figures 2C, D, and S1C). Finally, to assess the effect of cytosolic pH directly, we manipulated the cytosolic pH using the ionophore 2,4-dinitrophenol (DNP). Sch9 dissociated from the membrane in pH 6.5 buffer with DNP but not in pH 7.5 buffer (Figure 2E). Together, these findings suggest that cytosolic pH plays a regulatory role in Sch9 localization.

**Figure 2.**
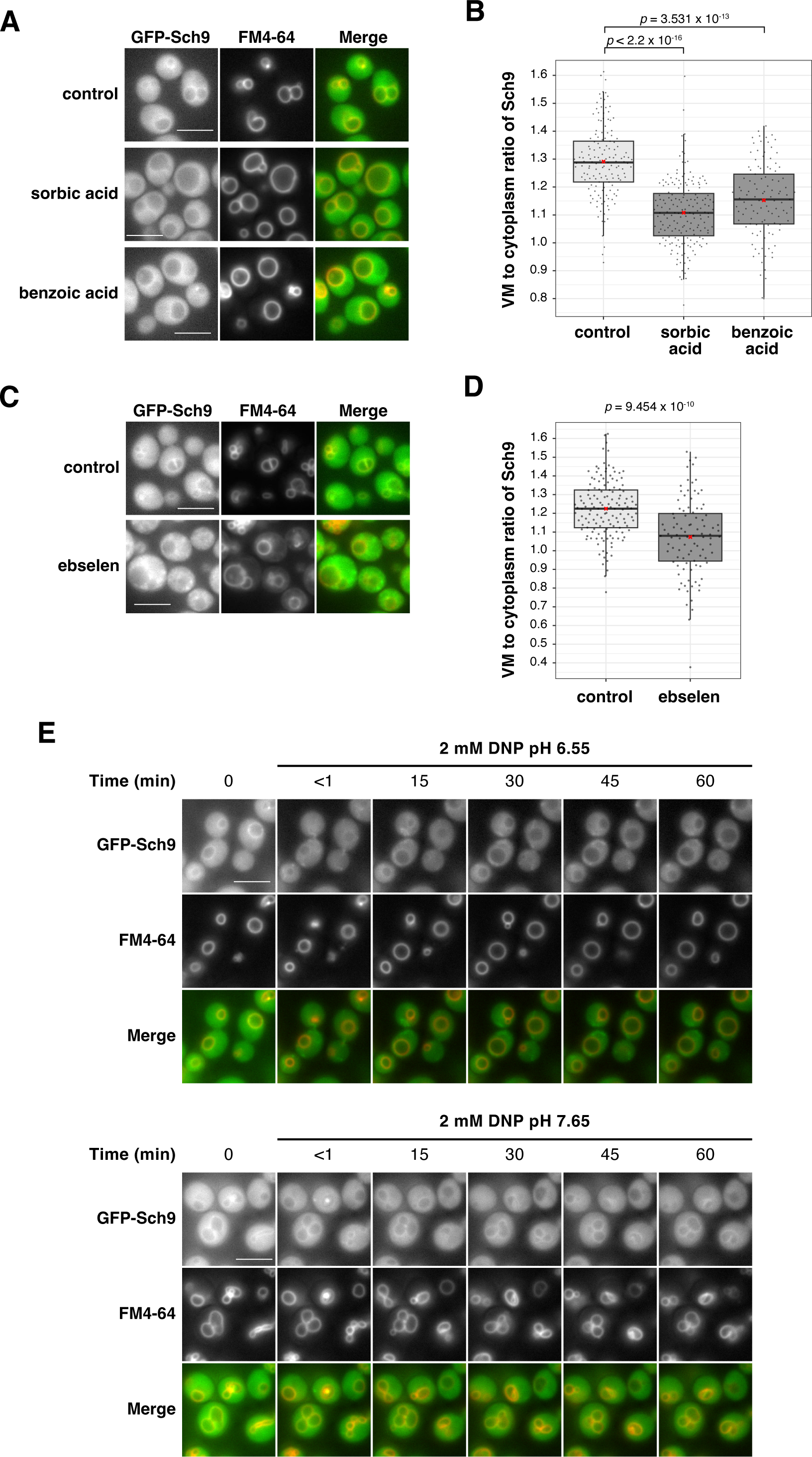
Sch9 dissociates from the vacuolar membrane in response to cytosolic acidification. **(A)** Fluorescent images of GFP–Sch9 and vacuole labeled by FM4–64. Exponentially growing cells were treated with 2 mM sorbic acid or 2 mM benzoic acid for 1 h. Control cells were incubated in SDCU medium for the same duration. **(B)** Quantitative analysis of the ratio of Sch9 fluorescence intensity at vacuolar membranes to that in the cytoplasm from (A). **(C)** Fluorescent images of GFP–Sch9 and vacuole labeled by FM4–64. Exponentially growing cells were treated with 100 μM ebselen for 5 min. Control cells were incubated in an SDCU medium for the same duration. **(D)** Quantitative analysis of the ratio of Sch9 fluorescence intensity of vacuolar membranes to that of the cytoplasm in (C). **(E)** Fluorescent images of GFP–Sch9 and a vacuole labeled with FM4–64. Exponentially growing cells were mounted onto a glass-bottom dish and treated with phosphate buffer adjusted to pH 6.55 or 7.65 containing 2 mM 2,4-dinitrophenol (DNP). Images were acquired before and at 15 min intervals after the initiation of DNP treatment. Box plots were constructed as described for Fig. 1. Scale bars = 5 μm. Significance was evaluated using the Steel test (B) and Brunner– Munzel test (D).

### Affinity of Sch9 and PI(3,5)P_2_ is pH-dependent

The protonation state of the phosphate headgroup in phosphoinositides containing PI(3,5)P_2_ is responsive to changes in pH (Kooijman et al., 2009), which could influence the affinity of lipid–protein interactions (Shin et al., 2020; Young et al., 2010). Therefore, we investigated whether the interaction between Sch9 and PI(3,5)P_2_ is pH-dependent. We conducted a liposome-binding assay using the purified recombinant N-terminal domain of Sch9 (Sch9^1–183^) and liposome containing 5% PI(3,5)P_2_ under varying pH conditions. Previous studies have demonstrated that Sch9^1–183^ is indispensable for binding to and sufficient for associating with the vacuolar membrane *in vivo* (Chen et al., 2021; Novarina et al., 2021). The binding of Sch9^1–183^ to the liposome decreased as the pH decreased (Figure 3A, B). This pH-dependent phenomenon was not observed when liposomes were used without PI(3,5)P_2_ (Figure S2). Thus, the interaction between Sch9 and PI(3,5)P_2_ appears to be pH-dependent, and this mechanism could explain the dissociation of Sch9 from the vacuolar membrane in response to cytosolic acidification.

**Figure 3.**
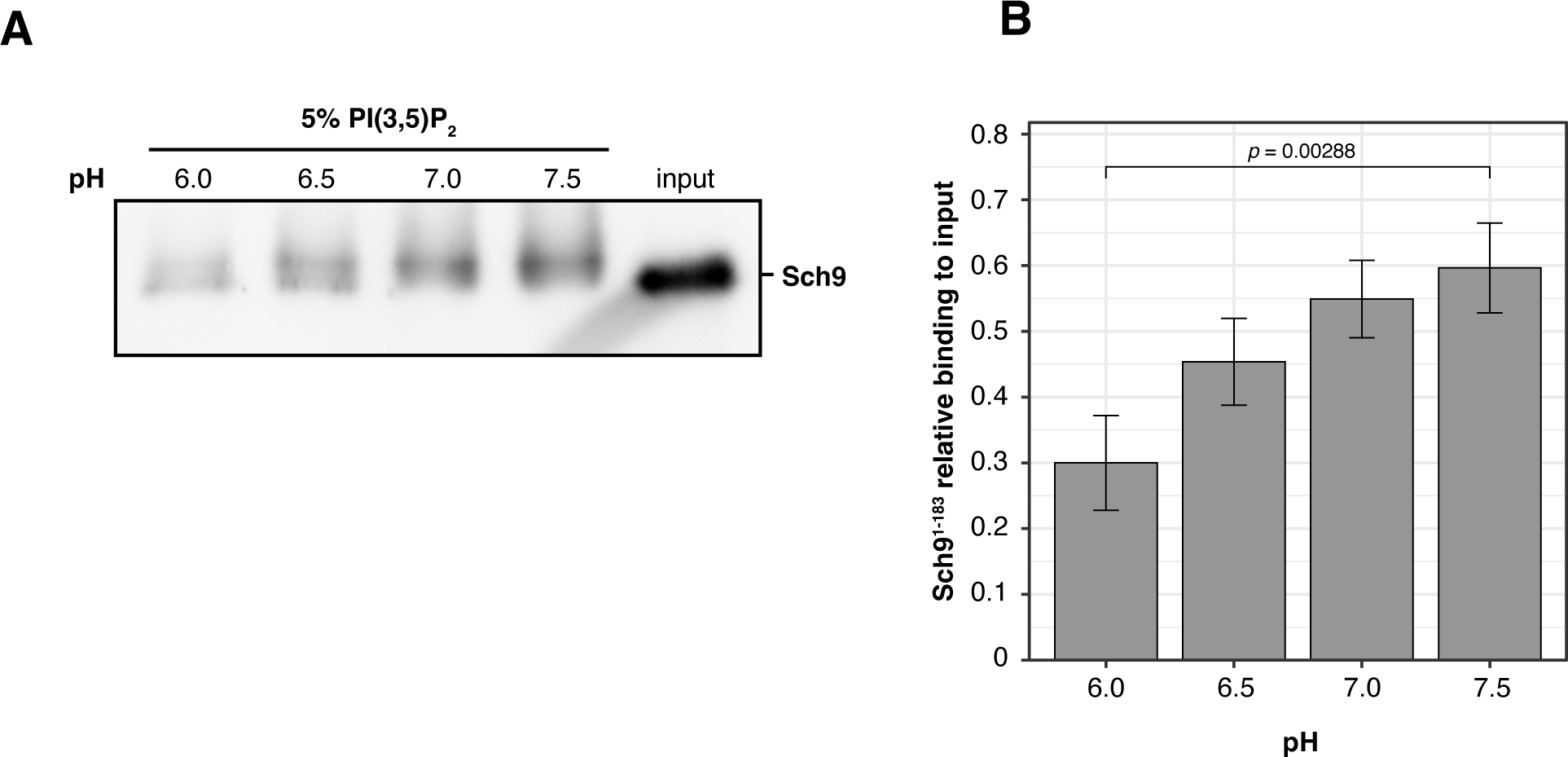
The affinity of Sch9 and PI(3,5)P_2_ is pH-dependent. **(A)** Western blotting analysis of recombinant Sch9^1–183^ binding to liposome containing 5% PI(3,5)P_2_ in buffers adjusted to various pH values (6.0–7.5). Input represents the total amount of protein used in the assay. **(B)** Quantitative analysis of the relative intensity to the input in (A). Data are mean ± standard error of the mean (SEM), based on three independent experiments. Significance was evaluated using Dunnett’s test, in which the relative intensity observed at pH 7.5 was compared to those at other pH levels.

### Dissociation of Sch9 from the vacuolar membrane promotes Sch9 dephosphorylation

The localization of Sch9 to the vacuolar membrane is required for TORC1-dependent phosphorylation (Novarina et al., 2021). Therefore, its pH-dependent dissociation could lead to dephosphorylation. To investigate this possibility, we constructed a strain in which Sch9 was artificially tethered to the vacuolar membrane through a mechanism independent of its binding to PI(3,5)P_2_. The strain was constructed to express a fusion protein of vacuolar membrane protein alkaline phosphatase Pho8 lacking the ALP activity (Pho8ΔALP) and GFP-binding protein (GBP), thereby ensuring the tethering of GFP–Sch9 to the vacuolar membrane via GFP–GBP interaction.

Initially, we confirmed that weak acid treatments did not lead to the dissociation of Pho8-tethered Sch9 from the membrane (Figure 4A). Subsequently, we compared the phosphorylation levels of WT Sch9 to that of Pho8-tethered Sch9 following weak acid treatment. Although the phosphorylation of WT Sch9 decreased, that of Pho8-tethered Sch9 remained unaffected by sorbic acid or benzoic acid treatment (Figures 4B, C, and S3A). This difference was specific to the TORC1–Sch9 pathway, as the phosphorylation of Pho8-tethered Sch9 is TORC1-dependent, and the phosphorylation state of Atg13 was not different between these strains (Figure S3B–D). Furthermore, at all tested concentrations, acetic acid caused less significant dephosphorylation in Pho8-tethered Sch9 than in the WT (Figure 4D, E). Unlike other weak acids, acetic acid treatment led to the restoration of Sch9 phosphorylation following its dephosphorylation. This could be attributed to the yeast’s robust capability to metabolize acetic acid and normalize cytoplasmic pH (Figure S3E). These findings indicate that the dissociation of Sch9 from the vacuolar membrane in response to a decrease in cytosolic pH contributes to Sch9 inactivation and selective regulation of the TORC1–Sch9 pathway.

**Figure 4.**
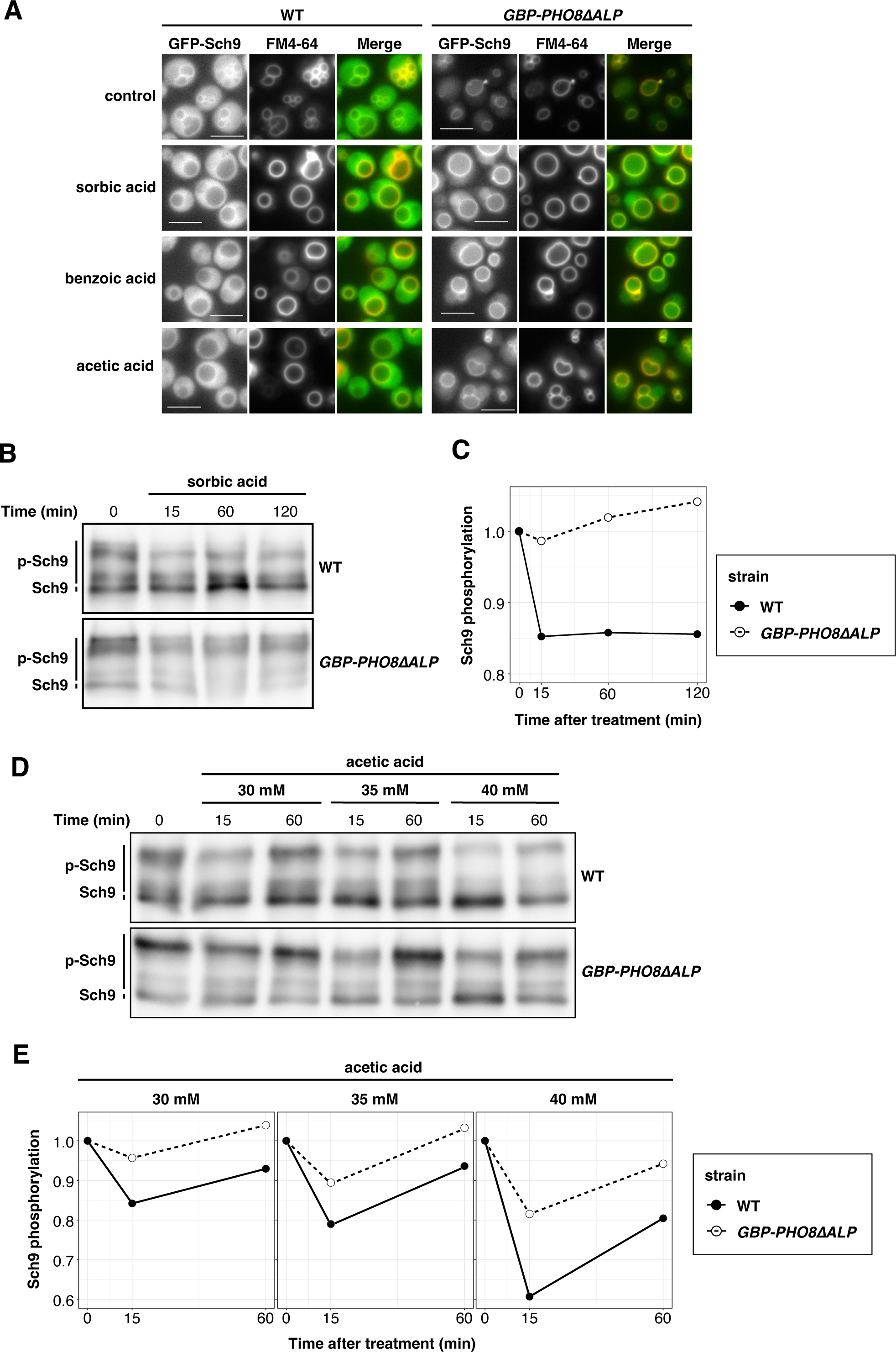
Dissociation of Sch9 from the vacuolar membrane promotes Sch9 dephosphorylation. **(A)** Fluorescent images of GFP–Sch9 and a vacuole labeled with FM4–64 of wild-type (WT) cells and cells expressing GBP–Pho8ΔALP. Exponentially growing cells were treated with 2 mM sorbic acid, 2 mM benzoic acid, or 35 mM acetic acid for 1 h. Control cells were incubated in SDCU medium for the same duration. Scale bars = 5 μm. **(B)** Western blotting analysis of the C-terminal fragment of Sch9–5HA in WT cells and cells expressing GBP– Pho8ΔALP. Exponentially growing cells were harvested before and at indicated time points following 2 mM sorbic acid treatment. The cell lysates were used in Western blotting. **(C)** Quantitative analysis of the band shift data shown in (B). **(D)** Western blotting analysis of the C-terminal fragment of Sch9–5HA in WT cells and cells expressing GBP–Pho8ΔALP. Exponentially growing cells were harvested before and at the indicated time points following treatment with acetic acid at concentrations of 30, 35, or 40 mM. **(E)** Quantitative analysis of the band shift data in (D).

### Detachment of Sch9 from the vacuolar membrane contributes to the regulation of PDS gene expression

In batch culture, medium glucose depletion induces metabolic reprogramming, in a process described as a diauxic shift. Sch9 is involved in this metabolic reprogramming through its role in the inhibitory regulation of Rim15 and Gis1 (Pedruzzi et al., 2003; Roosen et al., 2005). Because glucose depletion results in cytosolic acidification, we hypothesized that cytosolic pH acts as a secondary messenger for glucose and regulates the localization of Sch9, contributing to metabolic reprogramming. We examined whether changes in the localization of Sch9 to the vacuolar membrane corresponded with distinct growth phases. In exponentially growing cells, Sch9 was enriched at the membrane (Figure 5A, 4–10 h). By contrast, cells in a saturated culture exhibited dissociation (Figure 5A, 13–22 h). As expected, cytosolic pH was lower in cells in the saturated culture than in exponentially growing cells (Figure S4A). Consistent with the change in localization, the phosphorylation level of Sch9 diminished following the saturation of cell growth (Figure 5B).

**Figure 5.**
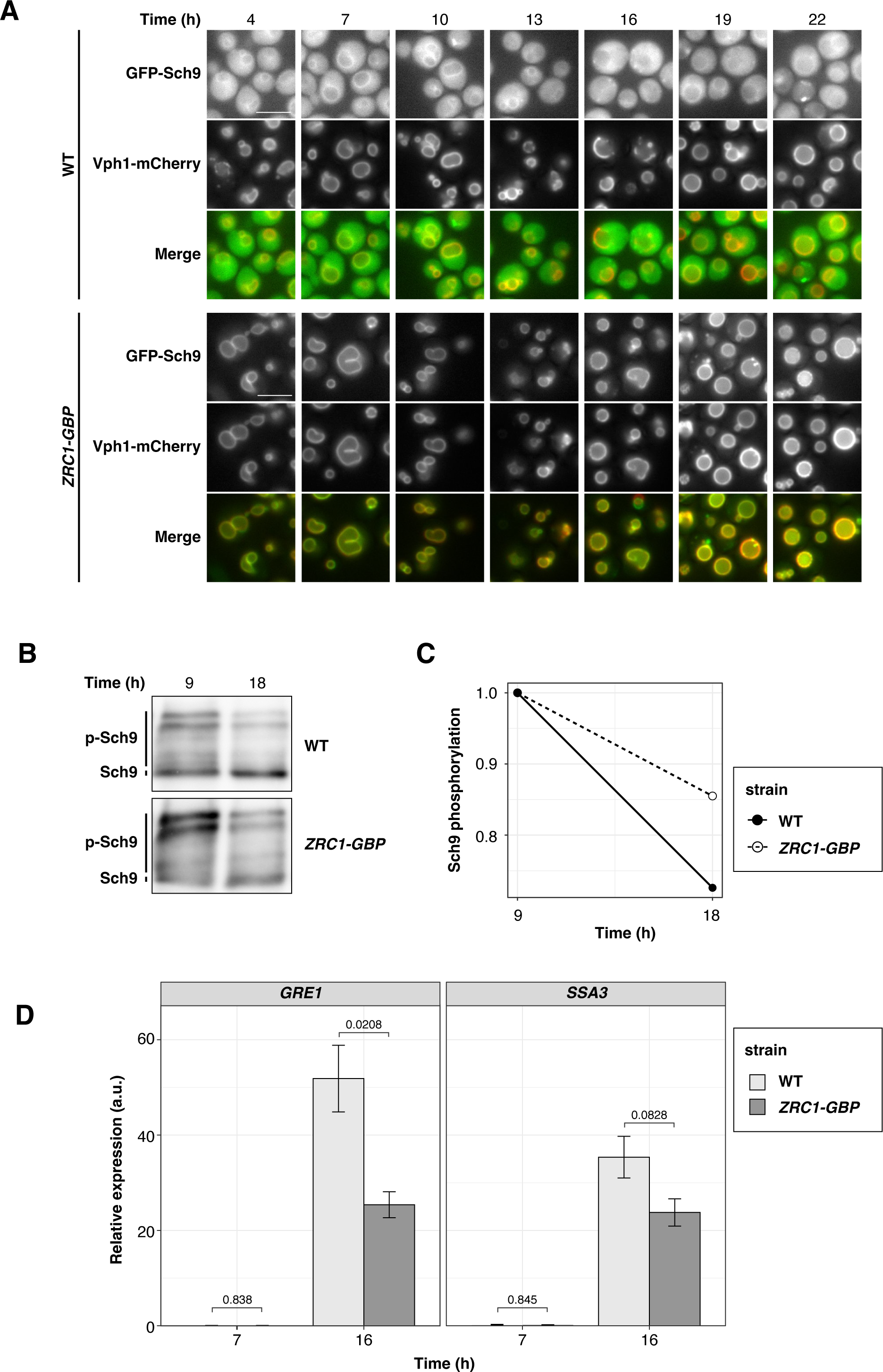
Detachment of Sch9 from the vacuolar membrane contributes to the regulation of post-diauxic shift (PDS) gene expression. **(A)** Fluorescent images of GFP–Sch9 and Vph1-mCherry of WT cells and cells expressing Zrc1-GBP. Overnight cultures were diluted into fresh SDCU medium at an optical density (OD_600_) of 0.1 (defined as time 0). Images were acquired at the indicated time points during cultivation. **(B)** Western blotting analysis of the C-terminal fragment of Sch9–5HA in WT cells and cells expressing Zrc1–GBP. Overnight cultures were diluted into fresh SDCU medium at an OD_600_ of 0.1 (time 0). Cells were harvested during cultivation at the indicated time points, and their lysates were used in Western blotting. **(C)** Quantitative analysis of the band shift data shown in (B). **(D)** Relative expression levels of *GRE1* and *SSA3* in cells during exponential growth (7 h) and after reaching growth saturation (16 h) in WT cells and cells expressing Zrc1–GBP. Overnight cultures were diluted into fresh SDCU medium at an OD_600_ of 0.1 (time 0). Cells were harvested at 7 and 16 h of cultivation, and total RNA was extracted. The mRNA levels were measured via quantitative reverse-transcription polymerase chain reaction (qRT-PCR). Data are means ± SEMs, based on three independent experiments. Significance was evaluated using Welch’s t-test.

To determine whether the inactivation of Sch9 could be attributed to its localization, we attempted to use a strain expressing GBP–Pho8ΔALP. However, due to the characteristics of Pho8 (He et al., 2021), GBP–Pho8ΔALP was incorporated into the vacuole as the glucose decreased. Therefore, instead of Pho8, we tagged GBP to Zrc1, a vacuolar membrane zinc transporter. We confirmed that this tagging did not influence cell growth (Figure S4B). In cells expressing Zrc1–GBP, Zrc1-tethered Sch9 persisted at high levels on the vacuolar membrane, even during the saturated growth phase (Figure 5A). The dephosphorylation of Zrc1-tethered Sch9 was attenuated compared to that of WT Sch9 (Figure 5B). Therefore, the dissociation of Sch9 from the vacuolar membrane associated with glucose consumption plays a role in the inactivation of the TORC1–Sch9 pathway.

To investigate the effect of changes in Sch9 localization on metabolic reprogramming, we measured the expression levels of PDS genes (*GRE1* and *SSA1*) in the WT strain and strain expressing Zrc1–GBP. The expression levels of the PDS genes were low, with no significant difference between the two strains during the glucose consumption phase (Figure 5C, 7 h). However, following cell growth saturation, induction of the PDS genes was suppressed in the strain expressing Zrc1–GBP compared to WT (Figure 5C, 16 h). Therefore, we conclude that the localization switch of Sch9 in response to cytosolic acidification upon glucose depletion promotes metabolic reprogramming, partially through regulating the expression of PDS genes.

### Dissociation of Sch9 from the vacuolar membrane confers an advantage for adaptation to acetic acid

The increase in extracellular acetic acid causes cytosolic acidification, leading to metabolic disturbance, the accumulation of reactive oxygen species (ROS), and mitochondrial dysfunction (Guaragnella and Bettiga, 2021). These cytotoxic effects inhibit cell proliferation, and higher concentrations of acetic acid induce programmed cell death (Chaves et al., 2021; Ullah et al., 2012). Lacking Sch9 confers resistance to acetic acid depending on Rim15 and Gis1 (Burtner et al., 2009). Thus, the dissociation of Sch9 from the vacuolar membrane could facilitate adaptation to acetic acid through the inactivation of Sch9, followed by the activation of Rim15 and Gis1. To test this possibility, we performed competition experiments between the same two strains (Figure 6A). Without acetic acid, the proportion of WT cells remained largely stable, whereas with acetic acid, it increased to ∼70% (Figure 6B). This indicates a competitive advantage of WT cells over cells expressing ZRC1–GBP of approximately 1.03-fold in the absence of acetic acid, which increased to approximately 1.35-fold in the presence of acetic acid (Figure 6C). For a more detailed analysis, we monitored the individual growth of each strain following exposure to acetic acid. The cells expressing Zrc1–GBP had extended lag times compared to WT cells rather than differences in growth rates during exponential growth or final yields (Figure 6D). Therefore, we conclude that the detachment of Sch9 from the vacuolar membrane could be advantageous for adaptation to acetic acid, as it appeared to aid the adaptation process during the lag phase.

**Figure 6.**
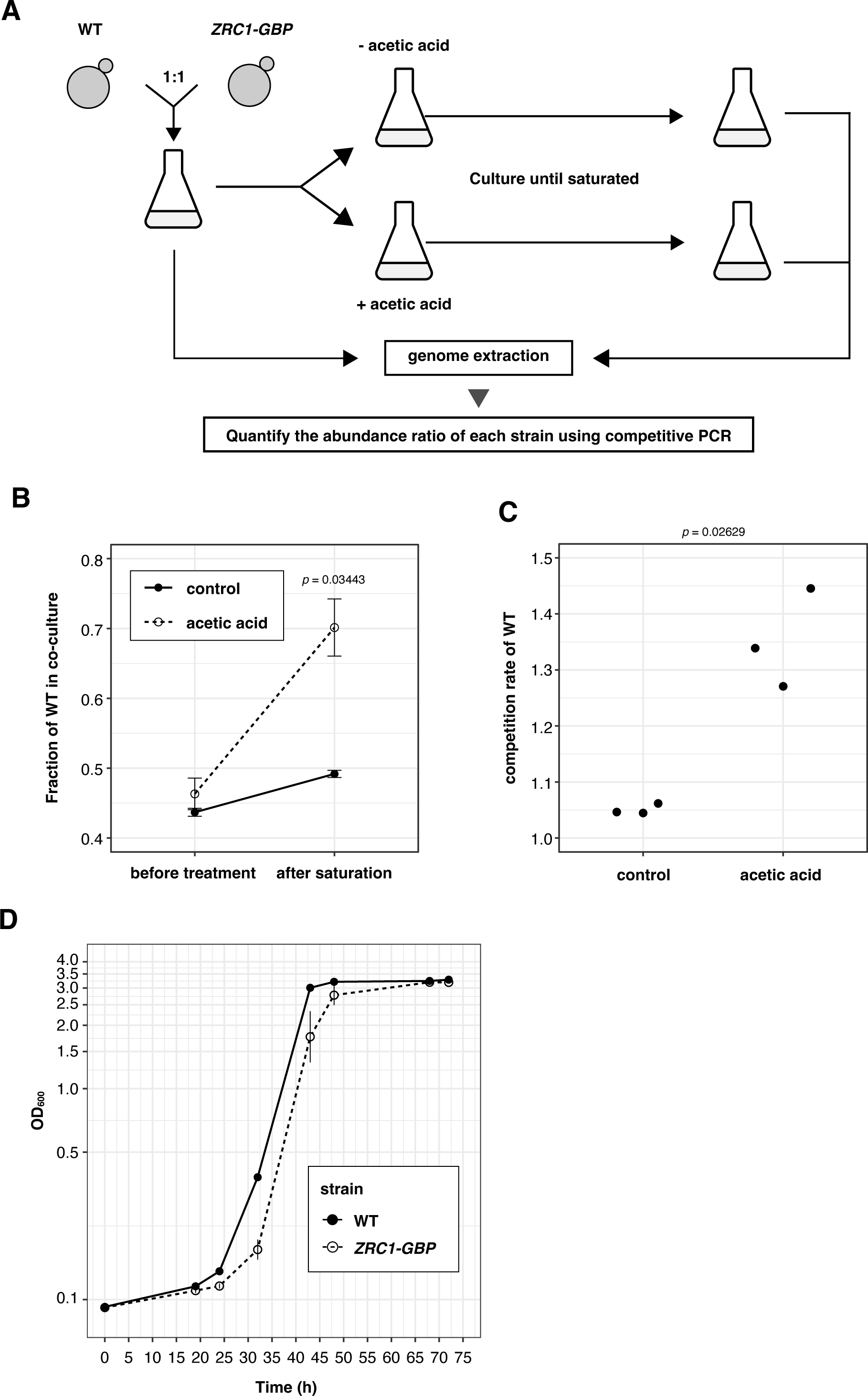
Dissociation of Sch9 from the vacuolar membrane confers an advantage for adaptation to acetic acid. **(A)** Schematic representation of the competitive growth assay. Detailed information is provided in the Materials and Methods section. **(B)** Fraction of WT cells in the competitive growth assay. WT cells and cells expressing ZRC1–GBP were mixed at an approximate 1:1 ratio and cultured in SDCU medium with or without 50 mM acetic acid until they reached growth saturation. The fraction of WT cells in cocultures was determined via competitive PCR. Data are means ± SEMs, based on three independent experiments. **(C)** Competition rate of WT cells calculated from (B). **(D)** Growth curves of WT cells and cells expressing Zrc1-GBP following acetic acid treatment. Exponentially growing cells were harvested, and the medium was replaced with SDCU medium containing 50 mM acetic acid at time 0. OD600 was measured at the indicated time points during cultivation. Data are means ± SEMs, based on three independent experiments. In (B) and (C), significance was evaluated using Welch’s t-test.

## Discussion

The regulation of anabolic and catabolic processes in response to environmental cues is essential for cellular survival. In budding yeast, the TORC1-Sch9 signaling pathway plays a pivotal role in this regulation. Previously, we demonstrated that changes in the localization of Sch9, driven by a mechanism that is not yet fully understood, is crucial for selectively regulating the TORC1–Sch9 pathway during oxidative stress (Takeda et al., 2017). In this study, we analyzed the mechanism of this localization switch. We observed localization under various stress conditions and presented lines of evidence demonstrating that the association between Sch9 and vacuolar membranes *in vivo* and the interaction between Sch9 and PI(3,5)P_2_ *in vitro* are pH-dependent. Thus, we propose that the localization of Sch9 functions as part of the cytosolic pH-sensing system.

### Various mechanisms can disturb cytosolic pH

The cytosolic pH is typically maintained around neutrality through strict regulation of the proton influx and efflux balance across the cellular membrane (Valkonen et al., 2013). Various stresses can disrupt this balance. For example, glucose depletion leads to rapid ATP depletion, thereby reducing cytosolic pH by impeding proton efflux from the cytosol, which is primarily mediated by the P-type ATPase Pma1 (Dechant et al., 2010). Weak acids can also contribute to lowering the cytosolic pH (Piper et al., 2001). When the extracellular pH is below their pKa, weak acids exist predominantly in a protonated form and can pass through the cellular plasma membrane more easily. Once inside the relatively neutral cytosol, these acids dissociate and release protons. Similarly, H_2_O_2_ has been observed to reduce cytosolic pH, although the exact mechanism underlying this process remains unclear (Dodd and Kralj, 2017). Two hypotheses are that hydroxyl radicals generated inside the cell may inhibit glycolysis, and/or H_2_O_2_ may act similar to a weak acid (Lin et al., 2018; Schalkwyk et al., 2013; Tsai et al., 1997; Zhao et al., 2019). Furthermore, factors that increase membrane permeability, such as heat shock, copper ions, and straight-chain alcohols, lead to increased proton influx, thereby lowering the cytosolic pH (Charoenbhakdi et al., 2016; Fernandes and Sá-Correia, 2001; Triandafillou et al., 2020).

Cytosolic acidification can inhibit various cellular functions such as enzyme activity, protein folding, and nutrient transport, and may also play a critical role in stress responses. For example, the drop in pH during glucose depletion triggers a cytoplasm transition from a liquid-like to a solid-like state, which aids cell survival under energy shortage conditions (Munder et al., 2016). In addition, transient cytosolic acidification during heat stress activates a transcriptional stress response facilitated by heat shock factor 1 (Hsf1) (Triandafillou et al., 2020). These observations suggest that cells have mechanisms to effectively utilize stress-induced pH reduction. The pH-responsive Sch9 localization may be part of this system.

### Changes in the protonation state of PI(3,5)P_2_ could explain its pH-dependent binding to Sch9

The protonation state of the phosphomonoester headgroup in phosphatidic acid (PA) and phosphoinositides varies within the physiological pH range and affects the affinity of protein–lipid interactions (Kooijman et al., 2009, 2005). Indeed, the interactions between Opi1 and PA, Osh1 and PI4P, and Vps34 and PI3P are modulated by pH (Naufer et al., 2017; Shin et al., 2020; Young et al., 2010). The electrostatic/hydrogen bond switch mechanism explains the pH dependency of protein–PA interactions (Kooijman et al., 2007). In this model, basic amino acid side chains are crucial for stabilizing protein interactions with PA. Sch9 contains four lysine and two arginine residues in its lipid-binding domain. These basic amino acid side chains might contribute to the pH-dependent affinity between Sch9 and PI(3,5)P_2_. Our experiments did not exclude the possibility that pH-sensitive changes in Sch9, rather than changes in the protonation state of PI(3,5)P_2_, function as pH sensors. Notably, the affinity of the FYVE domain for PI3P increases at lower pH levels, attributed to the protonation of two histidine residues within the FYVE domain due to reduced pH, increasing the positive charge (Lee et al., 2005). Due to the predominantly disordered structures in the N-terminal domain of Sch9, it remains challenging to predict its binding mode to PI(3,5)P_2_ and identify the amino acid residues involved in this interaction from a structural perspective (Caligaris et al., 2023).

There may be undiscovered protein–lipid combinations that show pH-dependent changes in these interactions. Comprehensive research is required to validate this hypothesis.

### Cytosolic pH selectively regulates TORC1–Sch9 signaling

Cytosolic pH is tightly regulated and affects cellular signaling pathways, including PKA, SNF, and TORC1 (Colombo et al., 1998; Dechant et al., 2014, 2010; Gutierrez et al., 2022; Isom et al., 2018; Simpson-Lavy and Kupiec, 2022; Thevelein, 1991). Our results are consistent with a previous study that found that the suppression of Pma1 expression using the tet-off system resulted in cytosolic acidification, leading to the dephosphorylation of Sch9 (Dechant et al., 2014). Moreover, FCCP protonophore treatment has been shown to induce cytosolic acidification and activate the TORC1–Npr1 pathway but not the TORC1–Sch9 pathway (Saliba et al., 2018). Notably, a recent study on the yeast phosphoproteome response revealed that the TORC1–Sch9 pathway specifically responds to pH perturbation, whereas the TORC1–Tap42–PP2A/Sit4 pathway does not (Leutert et al., 2023). These findings indicate that pH fluctuations impact the TORC1 pathway differently across various effectors and inhibit the TORC1–Sch9 pathway.

The downregulation of the TORC1–Sch9 pathway under diverse stress conditions highlights its role as a general stress response mechanism. Indeed, it is logical that biosynthetic processes would be halted during stresses to conserve cellular resources and energy. Given that decreased cytosolic pH is a common feature across multiple independent stressors, cytosolic pH may be useful as an integrative signal for these disparate stress conditions.

Therefore, detecting changes in cytosolic pH could be an effective mechanism for the TORC1–Sch9 pathway to respond comprehensively to a wide range of stressors.

### Sch9 localization on the vacuolar membrane facilitates the integration of TORC1–Sch9 signaling with various intracellular processes

Why is Sch9 abundant on the vacuolar membrane? First, our experiments demonstrated that it allows for the pH-dependent regulation of the TORC1–Sch9 pathway. This is mainly attributable to the localization of TORC1 itself at the vacuolar membrane. The presence of Sch9 close to TORC1 on the membrane probably facilitates efficient signal transduction.

Second, it may act as a sensor to monitor the state of the vacuole. The quantity of PI(3,5)P_2_ on the membrane can fluctuate depending on vacuole maturity and the cell cycle phase (Jin and Weisman, 2015; Okreglak et al., 2023). These dynamics may enable the coupling of vacuolar integrity and cell cycle progression with the activity of the TORC1–Sch9 pathway. Together, these findings indicate that the regulation of Sch9 via the amount of PI(3,5)P_2_ could play a role in the fine-tuning of Sch9 activity during cell proliferation, whereas pH-based regulation of Sch9 could be crucial for rapid responses to stresses.

The functional location of Sch9 may be significant. Given its known roles in the nucleus, Sch9 localization on the vacuolar membrane may be a first important step before sequestration by the nucleus (Jin et al., 2022; Oh et al., 2018; Pascual-Ahuir and Proft, 2007). The detachment of Sch9 from the vacuolar membrane may then facilitate its translocation into the nucleus, where it can function more effectively. In addition, Sch9 governs the assembly of the V-ATPase, and intriguingly, the pH-responsive Sch9 regulates the involvement of V-ATPase in pH regulation (Wilms et al., 2017). However, the specific effects of Sch9 location on V-ATPase assembly require further research to gain a more precise understanding of these potential roles.

### pH-dependent switching of Sch9 localization contributes to adaptation to acetic acid

Cells lacking Sch9 are more resistant to acetic acid, suggesting that Sch9 negatively regulates adaptation to acetic acid (Burtner et al., 2009; Deprez et al., 2021; Huang et al., 2012; Oh et al., 2018; Rego et al., 2020, 2018). Although further research is necessary to fully understand the precise mechanisms by which Sch9 inhibits the acetate adaptation process, several hypotheses have been proposed. First, the inactivation of Sch9 could promote tolerance to acetic acid by alleviating its repressive effects. Sch9 inhibits the activity of proteins and the expression of genes that are crucial for resistance to acetic acid. The expression of glucose-repressed genes plays a significant role in tolerance to acetic acid (Laera et al., 2016), and is facilitated by the transcription factors Adr1 and Cat8, whose expression is suppressed by Sch9 (Peterson and Liu, 2021). In addition, Sch9 inhibits Rim15 and Gis1, which are required for acetate tolerance (Burtner et al., 2009).

Second, Sch9 inactivation may help maintain functional mitochondria. Isc1, an inositol phosphosphingolipid phospholipase C, relocates from the endoplasmic reticulum (ER) to the mitochondria during acetic acid treatment in a Sch9-dependent manner (Rego et al., 2018). Such migration affects the mitochondrial sphingolipid balance, potentially increasing acetic acid sensitivity.

Third, Sch9 inactivation could lead to changes in gene expression that are advantageous under acetic acid stress through chromatin modifications. Sch9 promotes the phosphorylation of histone H3 threonine 11 (H3T11) during acetic acid exposure (Oh et al., 2018), which leads to acetic acid sensitivity through unknown mechanisms.

Our investigation revealed that the inability of Sch9 to detach from the vacuolar membrane leads to an extended lag phase before resuming proliferation after acetic acid treatment. This lag period allows for various adaptive responses to acetic acid, such as cytosolic pH stabilization (Ullah et al., 2012), inhibition of acetic acid influx (Mollapour and Piper, 2007), ROS removal (Guaragnella et al., 2019), and cell wall restructuring (Ribeiro et al., 2021).

Thus our findings indicate that such detachment may facilitate these adaptive processes through the abovementioned mechanisms or others yet to be identified. Moreover, considering the restoration of cytosolic pH in cells adapting to acetic acid and resuming proliferation (Ullah et al., 2012), the pH-responsive localization of Sch9 could offer a mechanism by which Sch9 activity is temporarily suppressed during acetate adaptation.

## Materials and Methods

### Yeast strains and plasmids

The yeast strains and plasmids used in this study are listed in Tables S1 and S2. All strains were derived from *S*. *cerevisiae* strain BY4741. Gene tagging was conducted using a conventional one-step polymerase chain reaction (PCR)-mediated method (Longtine et al., 1998). The primers used in this study are listed in Table S3.

To construct a strain expressing pHluorin2, yeast-codon-optimized pHluorin2 was amplified from pRF37 via PCR using the primers F1_pHluorin2 and tCYC1_pHluorin2. The PCR product was inserted into pRF34, which had been digested with SalI, using Gibson assembly. The resulting plasmid was used to amplify pTEF1–pHluorin2–tCYC1::HphMX by PCR with the primers HO-M13U F and HO-R1 R, and then the PCR product was transformed into the *HO* locus of BY4741.

### Growth conditions

Yeast cells were grown in SDCU medium consisting of 0.17% BD Difco yeast nitrogen base without amino acids or ammonium sulfate (BD Biosciences, Franklin Lakes, NJ, USA), 0.5% ammonium sulfate (Nacalai Tesque, Kyoto, Japan), 0.5% Bacto casamino acids (Thermo Fisher Scientific, Waltham, MA, USA), 2% glucose (Nacalai Tesque), and 20 μg/mL uracil (Fujifilm Wako Pure Chemical Corp., Richmond, VA, USA) at 30°C.

### Microscopy

Overnight yeast cell cultures (16 h at 30°C) were diluted into fresh SDCU medium at an optical density at 600 nm (OD_600_) of 0.05. Cells grown to the early log phase were collected via centrifugation and resuspended in either starvation medium or medium supplemented with specific reagents, depending on the experimental conditions. After incubation for the indicated time, the cells were collected again via centrifugation and observed using a DeltaVision imaging system (AppliedPrecision, Issaquah, WA, USA). For time-lapse imaging, exponentially growing cells were mounted onto a glass-bottom dish coated with Concanavalin A (Sigma-Aldrich, St. Louis, MO, USA). Images were acquired using Softworx image acquisition and analysis software. For FM4–64 staining, FM4–64 dye (Thermo Fisher Scientific) was added to log-phase cell cultures at a final concentration of 2 μM. After incubation for 1 h, the cells were washed with fresh medium and incubated for 1 h.

### Quantification of Sch9 localization

The localization of Sch9 on the vacuolar membrane was assessed by calculating the ratio of Sch9 present on the membrane to the total amount of Sch9 within the cytosol. The intensity of GFP–Sch9 within the membrane region, extracted from the FM4–64-stained image, was computed to determine localization. The intensity of GFP–Sch9 in the cytosol, defined as the entire cell region excluding the vacuolar region, was determined to represent Sch9 within the cytosol. To account for background fluorescence, the value of each region was adjusted by subtracting the autofluorescence value of that region when unlabeled cells were observed under the same conditions.

### Intracellular pH measurements

Intracellular pH was measured with the genetically encoded pH sensor pHluorin2 (Mahon, 2011). The pH of a calibration buffer (50 mM 2-morpholinoethanesulphonic acid [MES], 50 mM 4-(2-hydroxyethyl)-1-piperazineethanesulfonic acid [HEPES], 50 mM KCl, 50 mM NaCl, 200 mM ammonium acetate, 10 mM NaN_3_, and 10 mM 2-deoxyglucose) was adjusted using either NaOH or HCl to create a range of pH values from 4.5 to 8.5, at 0.5 pH intervals. Immediately before the calibration step, 15 mM monensin and 10 mM nigericin in EtOH were added to the buffer at final concentrations of 75 μM and 10 μM, respectively. Yeast cells were harvested via centrifugation and then resuspended in the buffer. After incubation for 15 min at room temperature, the cells were analyzed through flow cytometry on a CytoFLEX S flow cytometer (Beckman Coulter, Brea, CA, USA).

The calibration curves were generated using the following calculations:

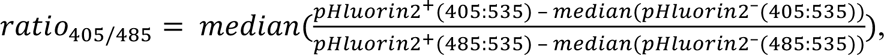

where pHluorin2^+^ represents pHluorin2-expressing cells and pHluorin2^−^ represents cells that do not express pHluorin2. Each point was fitted to the following sigmoid function:

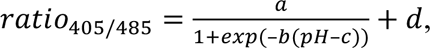

where a, b, c, and d are fitting parameters.

To measure cytosolic pH under various stress conditions, exponentially growing cells expressing pHlulorin2 or not were collected via centrifugation and resuspended in either starvation medium or medium supplemented with specific reagents, depending on the experimental conditions. After incubation for the indicated time, the cells were analyzed through flow cytometry. The cytosolic pH was determined by fitting the acquired ratio_405/485_ to the sigmoid function derived from the calibration curve.

In all conditions, cells expressing pHluorin2 or not were measured for background subtraction.

### Protein purification

Sch9 was expressed and purified from bacteria. To construct a plasmid harboring GST– Sch9^1–183^–6xHis, a DNA fragment containing Sch9^1–183^–6xHis, which was obtained via PCR amplification from the genomic DNA of BY4741 using the primers pGEX6P1_BamHI_SCH9_F and pGEX6P1_SCH9(1–183)_6His_R, was inserted into the pGEX-6P-1 vector digested with BamHI and EcoRI, using Gibson assembly. The obtained plasmid was introduced into the *Escherichia coli* strain BL21(DE3)pLysS. The bacteria were cultivated in 20 mL LB liquid media supplemented with 100 μg/mL ampicillin at 37°C overnight. Subsequently, the overnight culture was diluted in 750 mL LB media containing 100 μg/mL ampicillin to achieve an OD_600_ of 0.05 and cultured at 37°C until the OD_600_ was within the range 0.2–0.3. The cultivation temperature was lowered to 16°C, and the bacteria were further cultured until the OD_600_ reached 0.5. To induce expression of the target protein, a 0.9 M stock solution of isopropyl β-D-thiogalactopyranoside (IPTG) was added to the culture at a final concentration of 0.2 mM. The bacteria were cultured overnight, and then the bacterial cells were harvested and sonicated in cold phosphate-buffered saline (PBS). After centrifugation at 49,000 rpm for 30 min at 4°C, the supernatant was loaded onto a glutathione sepharose resin column. The column was washed with a wash buffer consisting of 50 mM Tris-HCl pH 8.0, 150 mM NaCl. Subsequently, 80 units of ProScission Protease were added to remove the GST tag, and the mixture was incubated at 4°C overnight. The eluate flowing out of the glutathione sepharose column was collected, and the purified protein was concentrated via ultrafiltration. The purified Sch9^1–183–^6xHis protein was stored in PBS with 10% glycerol at –30°C.

### Liposome binding assay

For the liposome binding assay, 5 µL of either 1 mM Control PolyPIPosomes or PI(3,5)P_2_ PolyPIPosomes (Echelon Biosciences, Salt Lake City, UT, USA) was rotated with 10 pmol purified Sch9^1–183–^6xHis in 250 µL liposome binding buffer (25 mM HEPES, 150 mM KCl, 25 μM CaCl_2_, 1 mM DTT) adjusted to the indicated pH using KOH for 30 min at room temperature. During incubation, Dynabeads M-280 Streptavidin (Thermo Fisher Scientific) was washed twice with liposome binding buffer. Following incubation, 20 µL beads was added to the mixture containing the liposomes and Sch9. The sample was rotated for 30 min at room temperature. Following incubation, a magnet was used to separate the beads from the supernatant. The supernatant was removed, and the beads were washed three times with 250 µL liposome binding buffer at the same pH. To avoid protein contamination from the tube wall, the resuspended beads were transferred to new tubes for each wash step. Then the beads were resuspended in 10 µL PBS and mixed with 10 µL 2× sample buffer (125 mM Tris-HCl pH 6.8, 4% sodium dodecyl sulfate, 20% glycerol, BPB, 10% 2-mercaptoethanol). After incubation for 10 min, the supernatant was separated from the beads using a magnet, boiled at 95°C for 5 min, and used for Western blotting analysis.

### Protein extraction and 2-nitro-5-thiocyanatobenzoic acid (NTCB) treatment

Protein extraction was performed as previously reported (Takeda et al., 2017). Briefly, cells in the early log phase or under stress conditions were treated with 6% trichloroacetic acid (TCA) and fixed for at least 30 min on ice. The cells were pelleted via centrifugation and washed twice with –30°C acetone. The pelleted cells were dried at room temperature. The pelleted cells were dissolved with 75 µL urea buffer (50 mM Tris-HCl pH 7.5, 6 M urea, 5 mM ethylenediaminetetraacetic acid [EDTA], 1% SDS, 5 mM NaF, 5 mM NaN_3_, 5 mM 4-nitrophenyl phosphate, 5 mM Na_4_P_2_O_7_, 5 mM β-glycerophosphate, 0.5× protease inhibitor cocktail (EDTA free) (Nacalai Tesque), 1.5 mM phenylmethylsulfonyl fluoride [PMSF]) and vortexed with the same volume of 0.5 mM low-alkaline glass beads for 10 min at 4°C. The cell lysates were heated at 65°C for 10 min and then centrifuged to separate soluble proteins. For cleavage of Sch9–5HA with NTCB (Tokyo Chemical Industry Co., Tokyo, Japan), the lysates were mixed with N-cyclohexyl-2-aminoethanesulfonic acid (CHES) buffer (pH 10.5) and NTCB in water at final concentrations of 100 mM and 1 mM, respectively, and incubated overnight at room temperature in the dark.

### Western blotting analysis

Protein lysates were boiled at 95°C for 5 min, loaded on an 8% SDS–polyacrylamide gel electrophoresis (PAGE) gel, and transferred to a polyvinylidene difluoride membrane (Merck, Rahway, NJ, USA). The membrane was incubated with primary antibodies overnight at 4°C. The membrane was washed and incubated with secondary antibodies for 60 min at room temperature. Immunoreactive proteins were visualized using ImmunoStar Zeta (Fujifilm Wako Pure Chemical Corp.). The protein abundance and band shift data were quantified using the Fiji distribution of the ImageJ software (National Institutes of Health, Bethesda, MD, USA).

### Antibodies

Sch9^1–183–^6xHis was detected using mouse monoclonal anti-His antibody (AB10597733, cat. # D291–3; MBL International, Woburn, MA, USA). A C-terminal fragment of Sch9–5HA was detected using a mouse monoclonal anti-HA antibody (AB2565007, cat. # 901502; BioLegend, San Diego, CA, USA). Atg13 was detected using an anti-Atg13 antibody gifted by Dr. Ohsumi (Tokyo Institute of Technology, Tokyo, Japan).

### Quantitative reverse-transcription PCR (qRT-PCR)

Total RNA was extracted using a hot acid phenol with glass bead extraction. Briefly, 3 (log phase) or 8 (stationary phase) OD_600_ units of cells were harvested, pelleted via centrifugation, and quickly frozen with liquid nitrogen. The cell pellets were resuspended in 270 µL AE buffer (50 mM sodium acetate pH 5.3, 10 mM EDTA pH 8.0) and 30 µL 10% SDS. Subsequently, approximately 0.3 g 0.5 mM low-alkaline glass beads and 300 µL preheated phenol saturated with citrate buffer at 65°C were added. The mixture was vortexed for 1 min and incubated at 65°C for 5 min. This cycle was repeated six times in total. The resulting cell lysate was subjected to standard pheno-chloroform extraction and ethanol precipitation. The extracted total RNA was reverse-transcribed into cDNA using ReverTra Ace qPCR RT Master Mix with gDNA Remover (Toyobo, Osaka, Japan). Real-time PCR was performed using THUNDERBIRD Next SYBR qPCR Mix (Toyobo) with three biological replicates, each consisting of two technical replicates. Relative mRNA levels were calculated using the ΔCt method, with the geometric mean of expression levels for UBC6, TAF10, and ALG9 utilized for normalization.

### Competitive growth assay

Exponentially growing WT cells and cells expressing Zrc1–GBP were diluted to an OD_600_ of 0.1 and mixed at equal volumes. Immediately, a sample of this mixture (optical density unit = 1) was collected, pelleted, and subsequently frozen (t_0_). The remaining cells underwent centrifugation and were resuspended in SDCU medium, with or without 50 mM acetic acid. Then these cells were incubated at 30°C until reaching saturation. After incubation, the cells were collected, pelleted, and frozen (t_1_). Genomic DNA was extracted from these pellets, and competitive PCR was performed using primers ZRC1 checkF and ZRC1 checkR. The ratio of PCR product from each strain was quantified. To create a calibration curve depicting the relationship between these PCR product ratios and the corresponding cell ratios, genomic DNA was extracted from cell mixtures prepared at defined ratios (WT:ZRC1–GFP = 1:4 to 4:1). Then this DNA was subjected to the same competitive PCR procedure. The resultant standard curve was utilized to determine the percentage composition of each cell line in the mixtures.

The competition rate of WT strains was defined as follows:

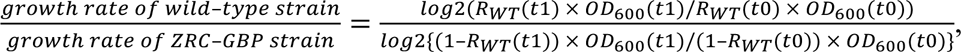

where R_WT_(t_1_) and R_WT_(t_0_) represent the proportions of WT cells at times t_1_ and t_0_, respectively. OD_600_(t_1_) and OD_600_(t_0_) represent the optical densities of the coculture at times t_1_ and t_0_, respectively.

## Acknowledgments

We thank the members of the Matsuura laboratory for valuable discussions, and Textcheck for editing the manuscript. This work was supported by JST SPRING, Grant Number JPMJSP2109, and a research grant from Institute for Fermentation, Osaka, Japan.

---

**Figure S1.**
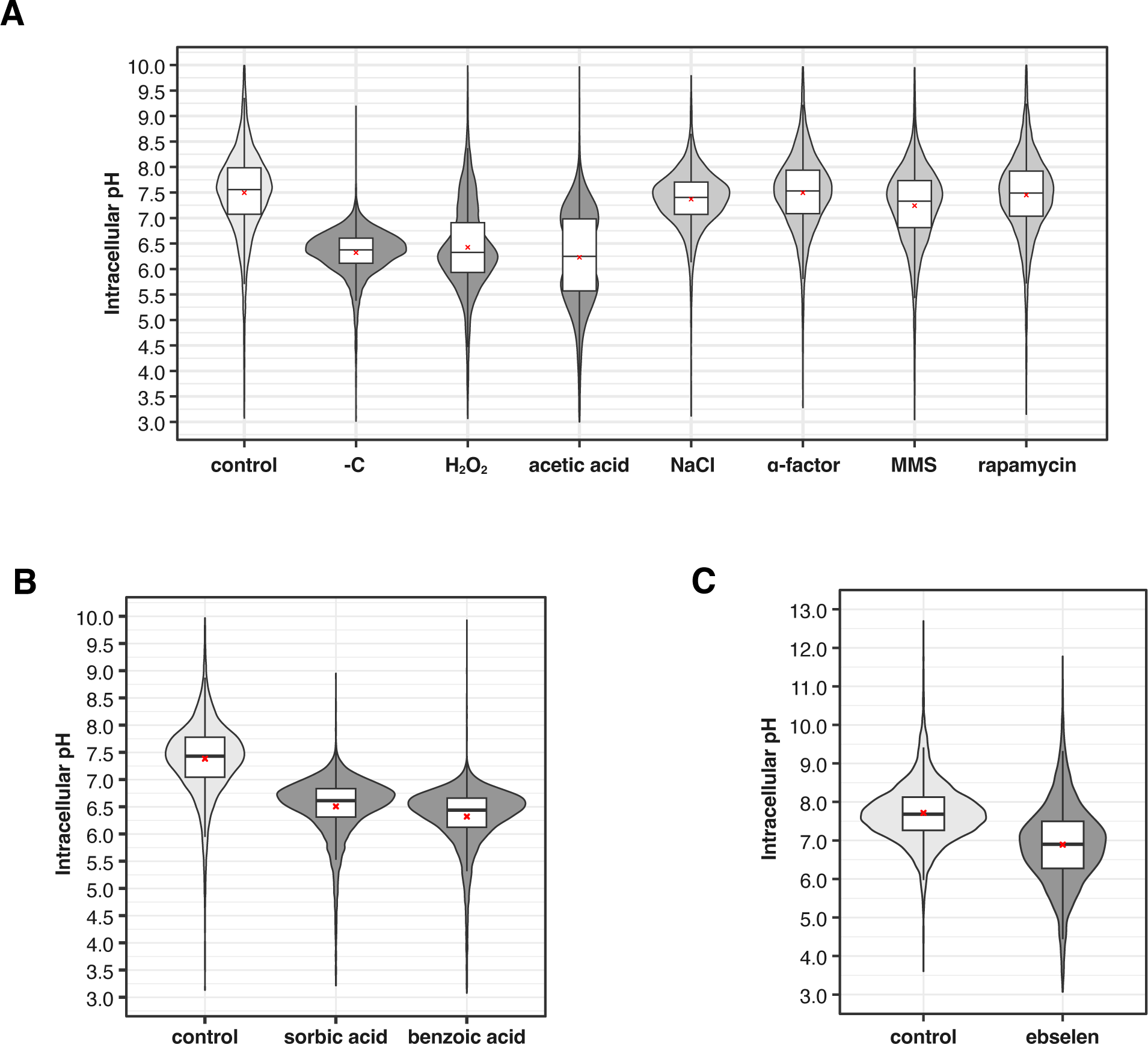
Cytosolic pH under various conditions. Distributions of cytosolic pH under identical stress conditions as described for Fig. 1 **(A)**, Fig. 2A **(B)**, and Fig. 2C **(C)**. The cytosolic pH of individual cells was determined by measuring pHluorin2 through flow cytometry. Box plots were constructed as described for Fig. 1.

**Figure S2.**
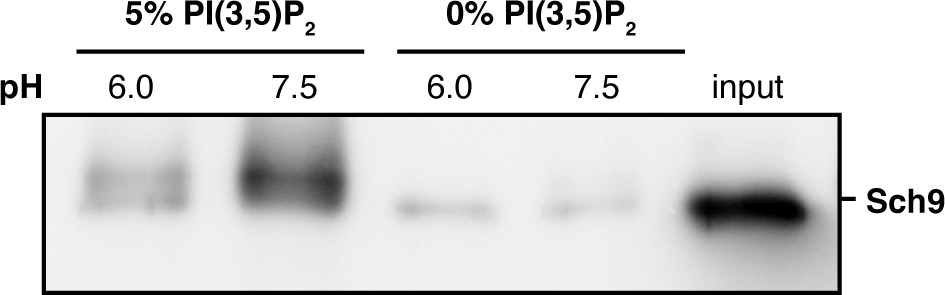
Binding of Sch9 to liposomes depends on PI(3,5)P_2_. Western blotting analysis of recombinant Sch9^1–183^ bound to liposomes with or without 5% PI(3,5)P_2_ in buffers adjusted to pH 6.0 or 7.5. Input represents the total amount of protein used in the assay.

**Figure S3.**
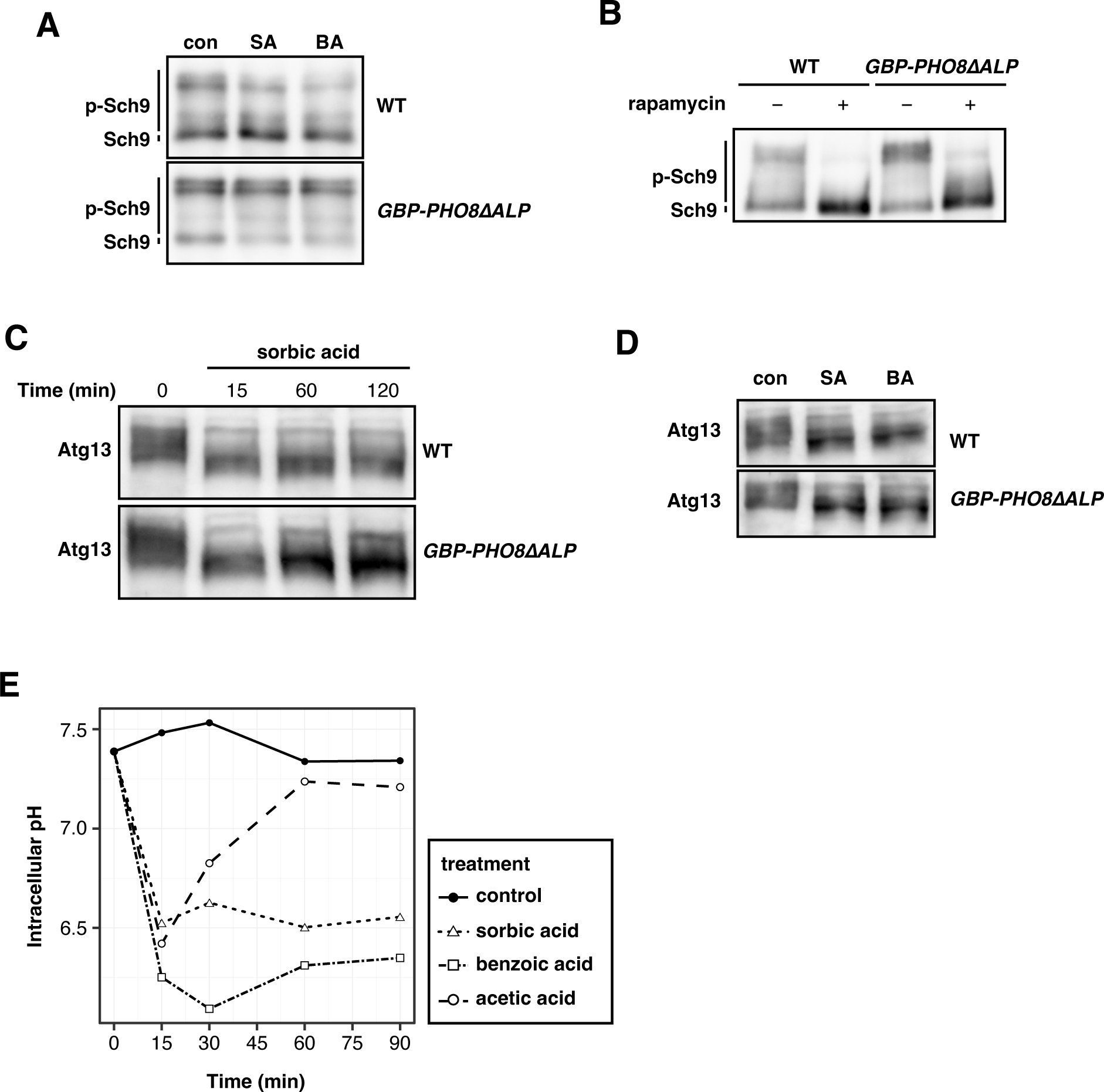
Dissociation of Sch9 from the vacuolar membrane contributes to the selective regulation of the TORC1–Sch9 pathway. **(A)** Western blotting analysis of C-terminal fragment of Sch9–5HA in WT cells and cells expressing GBP-Pho8ΔALP. Exponentially growing cells were harvested before and after treatments with 2 mM sorbic acid or 2 mM benzoic acid for 1 h. The cell lysates were used in Western blotting. **(B)** Western blotting analysis of C-terminal fragment of Sch9–5HA in WT cells and cells expressing GBP–Pho8ΔALP. Exponentially growing cells were harvested before and after treatment with 200 ng/mL rapamycin for 30 min. The cell lysates were used in Western blotting. **(C)** Western blotting analysis of Atg13 in WT or cells expressing GBP–Pho8ΔALP. The cell lysates used in this analysis were the same as those used in Figure 4B. **(D)** Western blotting analysis of Atg13 in WT or cells expressing GBP–Pho8ΔALP. The cell lysates used in this analysis were the same as those used in (A). **(E)** Cytosolic pH changes during weak acid stress treatment measured by pHluorin2. Exponentially growing cells were treated with 2 mM sorbic acid, 2 mM benzoic acid, or 35 mM acetic acid. Control cells were incubated in SDCU medium. Each point represents the average value of pH for all cells at each time point.

**Figure S4.**
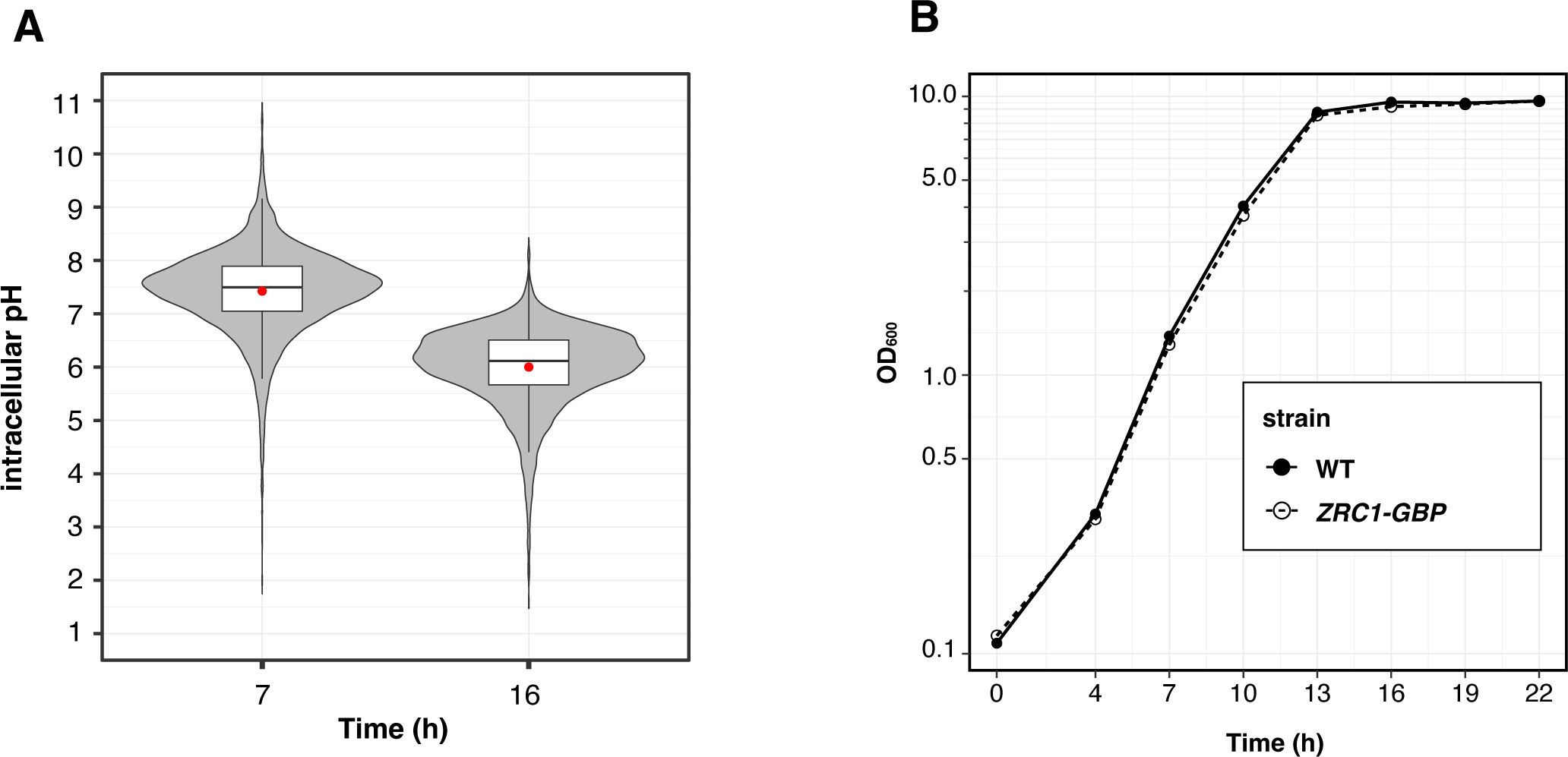
Cytosolic pH decreases following the saturation of cell growth. **(A)** Distributions of cytosolic pH in cells during exponential growth (7 h) and after reaching growth saturation (16 h). The cytosolic pH of individual cells was determined by measuring pHluorin2 through flow cytometry. Red crosses indicate mean values. **(B)** Growth curves of WT cells and cells expressing Zrc1–GBP. Overnight cultures were diluted into fresh SDCU medium at an OD_600_ of 0.1 (time 0). OD_600_ was acquired at the indicated time points during cultivation.

**Table S1.**
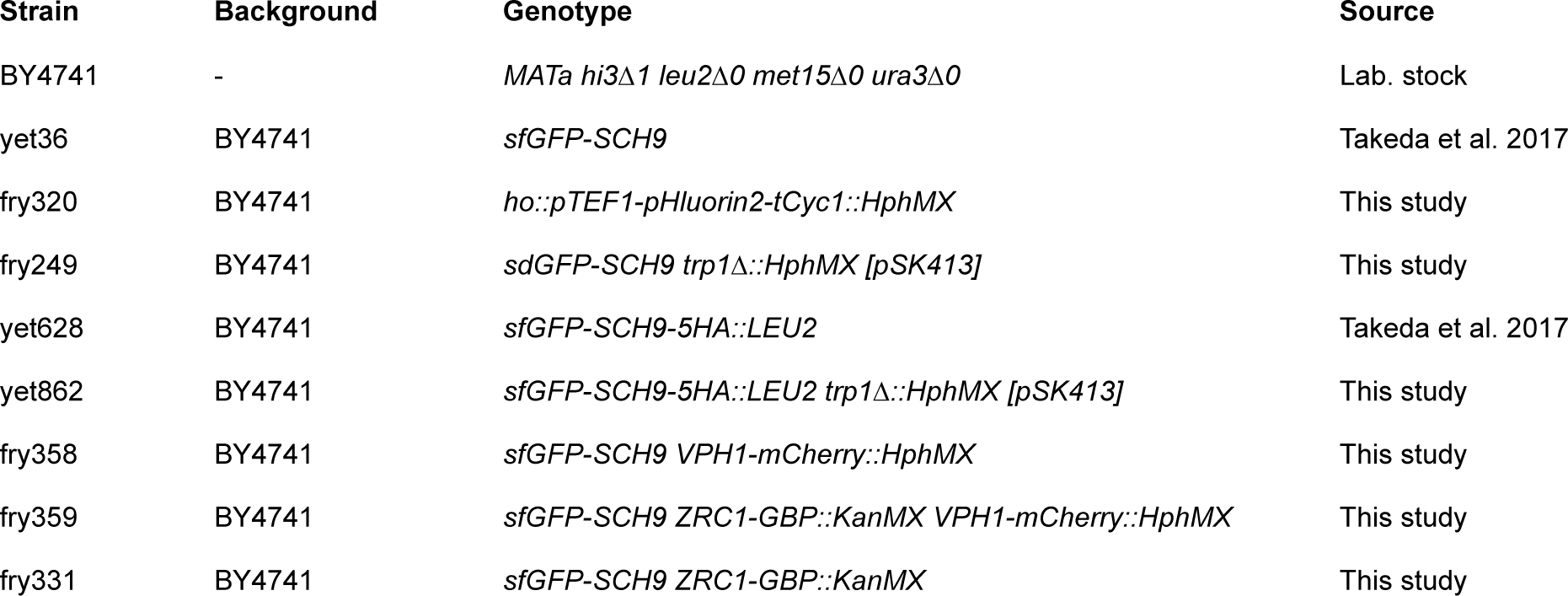
A list of yeast strains.

**Table S2.**
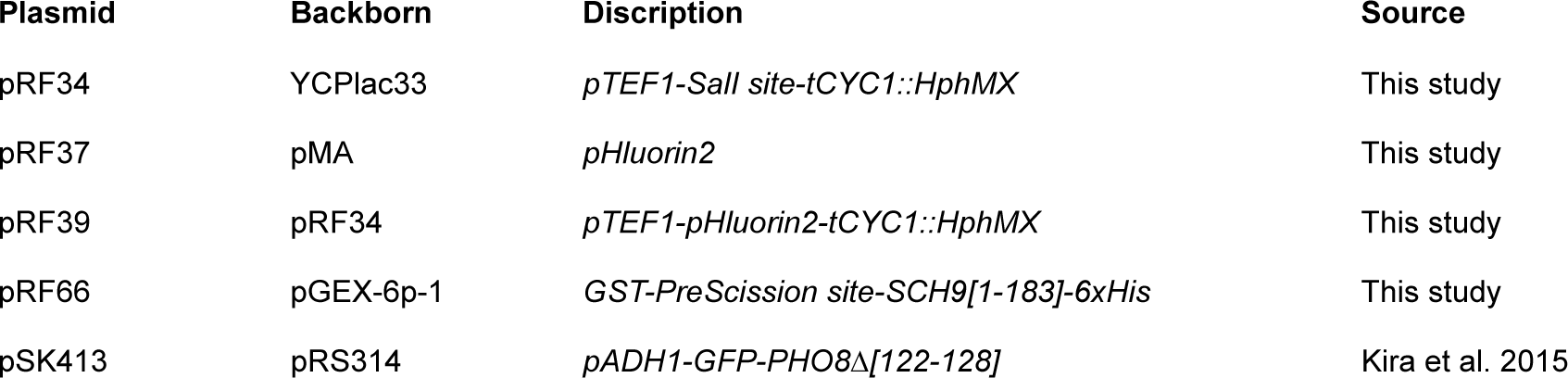
A list of plasmids.

| **Plasmid** | **Backborn** | **Discription** | **Source** |
| --- | --- | --- | --- |
| pRF34 | YCPlac33 | *pTEF1-Sall site-tCYC1::HphMX* | This study |
| pRF37 | pMA | *pHluorin2* | This study |
| pRF39 | pRF34 | *pTEF1-pHluorin2-tCYC1::HphMX* | This study |
| pRF66 | pGEX-6p-1 | *GST-PreScission site-SCH9[1-183]-6xHis* | This study |
| pSK413 | pRS314 | *pADH1-GFP-PHO8Δ[122-128]* | Kira et al. 2015 |

**Table S3.**
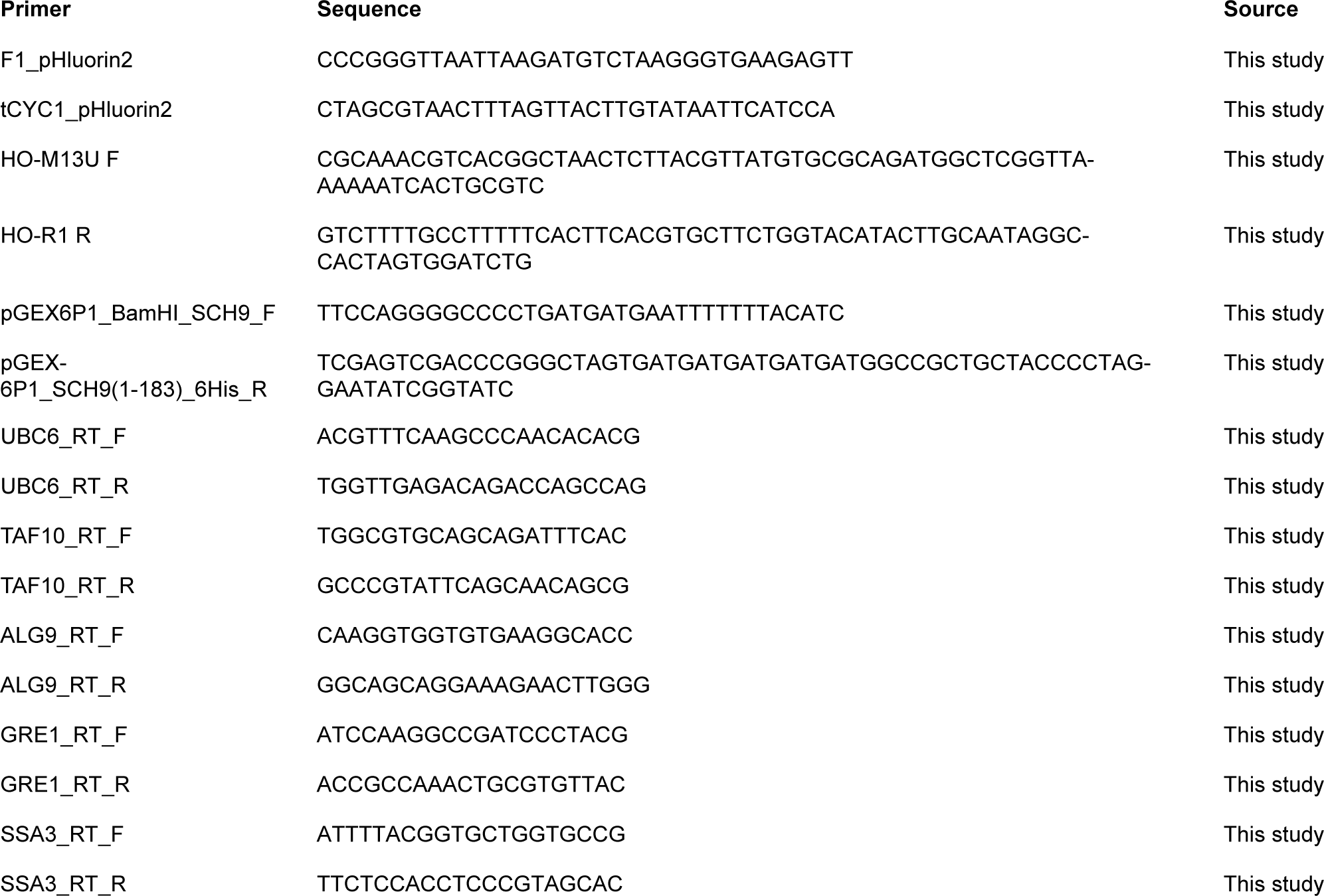
A list of primers.

